# MSCypher: an integrated database searching and machine learning workflow for multiplexed proteomics

**DOI:** 10.1101/397257

**Authors:** Eugene A. Kapp, Giuseppe Infusini, Yunshan Zhong, Laura F. Dagley, Terence P. Speed, Andrew I. Webb

## Abstract

Improvements in shotgun proteomics approaches are hampered by increases in multiplexed (chimeric) spectra, as improvements in peak capacity, sensitivity or dynamic range all increase the number of co-eluting peptides. This results in diminishing returns using traditional search algorithms, as co-fragmented spectra are known to decrease identification rates. Here we describe MSCypher, a freely available software suite that enables an extensible workflow including a hybrid supervised machine learned strategy that dynamically adjusts to individual datasets. This results in improved identification rates and quantification of low-abundant peptides and proteins. In addition, the integration of peptide *de novo* sequencing and database searching enables an unbiased view of variants and high-intensity unassigned peptide spectral matches.

**Highlights:**

- Open-source end-to-end label-free proteomics workflow
- Integrated database searching and machine learning
- Customisable and extensible workflow including *de novo* sequencing
- Optimised for multiplexed spectra, challenging proteomics datasets and peptidomics applications

## Main

Increased sensitivity of the latest generation mass spectrometers, in particular the UHR- QTOFs, has improved the measurable intra-scan dynamic range in which accurate peptide masses can be detected. Current instruments are capable of detecting >230,000 individual peptide isotopic patterns in a single 90 minute run (Beck et al., 2015), yet only a small fraction (10-20%) of these are identified (Chapman et al., 2014; Houel et al., 2010; Wang et al., 2011). Co-eluting peptide features are now able to be detected in over a 100,000-fold quantitative range, resulting in a drastic improvement in number of overall features detected. For a single MS run, a total of 235,389 isotope clusters can be detected using standard criteria (Cox and Mann, 2008) as shown in **Figure 1a**. A standard data-dependent approach clearly demonstrates the inherent under-sampling from complex biological mixtures, where only ∼22% of the peptide features were targeted for fragmentation and ∼10% of these were identified. From a 90 minute LC/MS run of a standard tryptic digest containing human and *E. coli* proteomes (600 ng), chimeric spectra accounted for more than 88% of the acquired data (**Figure 1b**) likely causing suppression of identification rates (Houel et al., 2010). While this has generally been thought detrimental to data-dependent acquisition (DDA) identification performance (Shishkova et al., 2016), this natural multiplexing provides an opportunity to increase identification rates. Attempts to utilize chimeric spectra was first described in 2000 (Masselon et al., 2000) and more recent attempts have proven quite successful (Cox et al., 2011; Shteynberg et al., 2015; Wang et al., 2011; Zhang et al., 2014; Zhang et al., 2005). However, these approaches all utilize the same peptide search strategies that are based on rational probabilistic scoring functions that only take into account a small fraction of information available to describe a potential peptide spectral match (PSM). Recent increases in computational power and the corresponding access to high quality training data has facilitated the ubiquitous use of machine learning for increased depth of data mining and this concept was first pioneered in the proteomics field by Kåll *et al.* (Kall et al., 2007). Indeed, post-processing of search results using more information in combination with a semi-supervised machine learning approach (Granholm et al., 2014) can drastically improve peptide identification rates. We now propose to extend this idea to a learned matching classification strategy which classifies target PSM from decoy PSM, incorporating as many parameters as possible to describe matching (parameters listed in **Table S1**). MSCypher is a hybrid workflow (**Figure 1c**) that currently utilizes the feature detection from the MaxQuant workflow and consists of a combined pre-matching and sensitive search algorithm that interfaces with a supervised machine learning classification using the random forest algorithm (Breiman, 2001). In testing early implementations, we used Monte-Carlo cross validation (MCCV) to calculate accuracy, true positive and false positive rates (TPR, FPR) and their confidence intervals (**Figure S1**). We observed that varying sizes of randomly chosen training/test sets, the classification model consistently generated accuracies in excess of 99.5%. When comparing scoring outputs of different search algorithms, the RF-generated classification score exhibited a wider separation between potentially correct identifications and decoy matches (**Figure S2)**. The current implementation of the workflow incorporates existing approaches including feature detection, first-pass searching for *m/z* and retention time (RT) re-calibration, peak picking (Cox and Mann, 2008) and RT prediction (Moruz et al., 2012).

**Figure 1.**
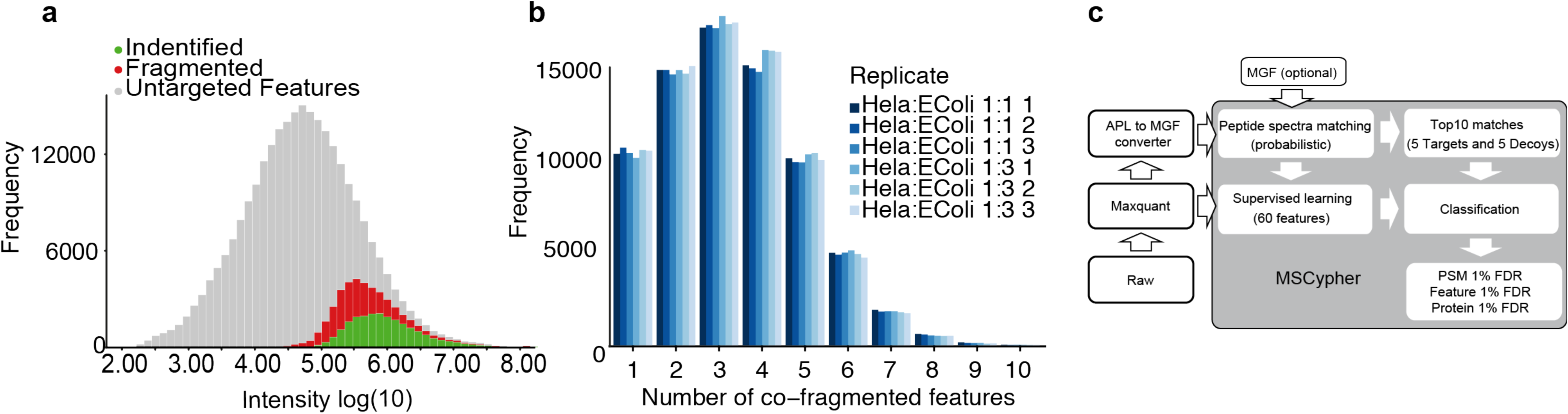
(a) Histogram of frequency of the detected features binned by intensity during a single LC/MS run. Un-fragmented isotopic features (gray), fragmented features (red) and identified isotopic features (green) demonstrate the small fraction of the >230,000 detected peptide features identified in current proteomics workflows. (b) Histogram representing frequency of co-fragmented peptide features within each isolation window. (c) MSCypher workflow.

To compare the effect of highly chimeric spectra on matching algorithms, we simulated datasets to measure multiplexed identification rates, first proposed in 2000 (Masselon et al., 2000). From a consensus set of 80,000 peptide spectra identified by Mascot, MSGF+, Comet and MSCypher, we iteratively selected 100 identified spectra at random. To simulate increasing chimericity, at each iteration, we appended from 1 to 10 additional MS/MS spectra selected at random to the peak list. This was repeated 10,000 times to generate a total of 1 million randomly generated chimeric spectrum entries per iteration. We then compared the ability of MSCypher with both the machine learning step switched on and off, to correctly re-call the consensus primary sequence (**Figure 2a**). The highest fraction of correctly called spectra across all iterations was obtained by including machine learning (ML). While the underlying probabilistic peak matching from MSCypher performed strongly as well, the increase in correct peptide calls when using ML represent spectra that have been re-ranked from lower scoring matches. This ability of the classification approach to re-rank potential matches represents a unique ability among peptide-based search algorithms as this represents evidence that has not been published to-date using post-scoring machine learning algorithms.

**Figure 2.**
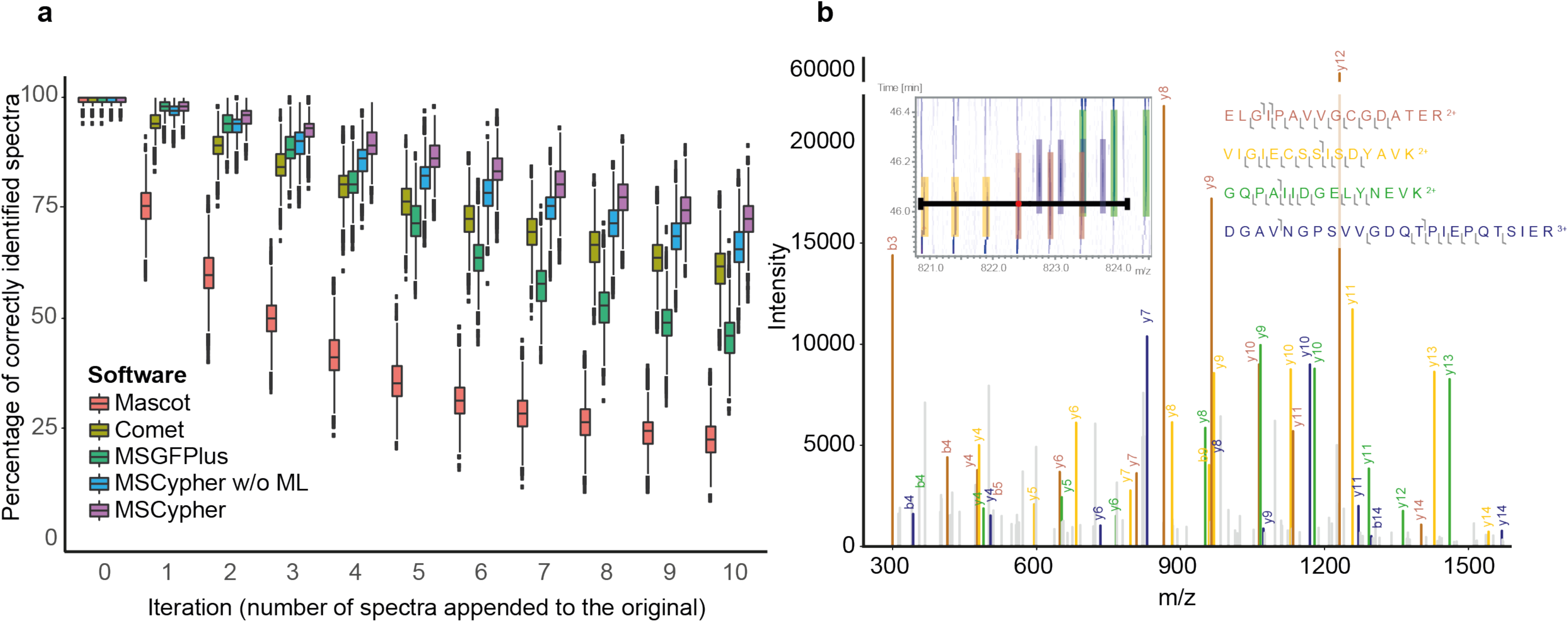
Comparison of search engines using simulated chimeric spectra. (a) Comparison of common search algorithms using a Monte Carlo simulation of increasing chimeric spectral complexity (b) Example of observed chimeric spectra obtained from co-fragmentation of four peptide features, one being the picked precursor ion (red), second (yellow), third (green) and fourth (blue). Matching y and b-ions are colored according to precursor. Inset shows the 2D LCMS region of interest, where the black line represents the isolation window and the colored bars the detected isotopes of the co-fragmented precursors.

To demonstrate the improvements that can be gained from ML, we analysed a tryptic digest of human HeLa cell lysate that was seeded with *E. coli* peptides. Here we observed four separate peptide features identified from a single isolation and fragmentation event (**Figure 2b inset**). The spectra present as a typical MS/MS spectrum and four peptides were identified below a 1% FDR using the MSCypher workflow (**Figure 2b**).

As high-quality feature detection is a fundamental requirement of accurate database searching, the current implementation of MSCypher requires first processing raw data using the MaxQuant program. Thus, the widely-used Andromeda search engine (Cox et al., 2011) is a natural search algorithm with which to compare and benchmark our software. MSCypher consistently yielded higher PSMs at 1% filtered false discovered rate (FDR) for all replicate runs, including both instrument picked precursors and secondarily identified spectra from co-fragmented precursors (**Figure 3a**). Interestingly, comparing the precursor density distribution of identified peptides from each algorithm with respect to the peptide intensity (**Figure 3b**), reveals that the additional identifications are not correlated to precursor intensity, but appear to be completely new matches that MaxQuant did not identify. This is reflected in the increased total peptide sequence count (**Figure 3c**) and resulted in increased sequence coverage of the majority of inferred proteins (**Figure 3d**).

**Figure 3.**
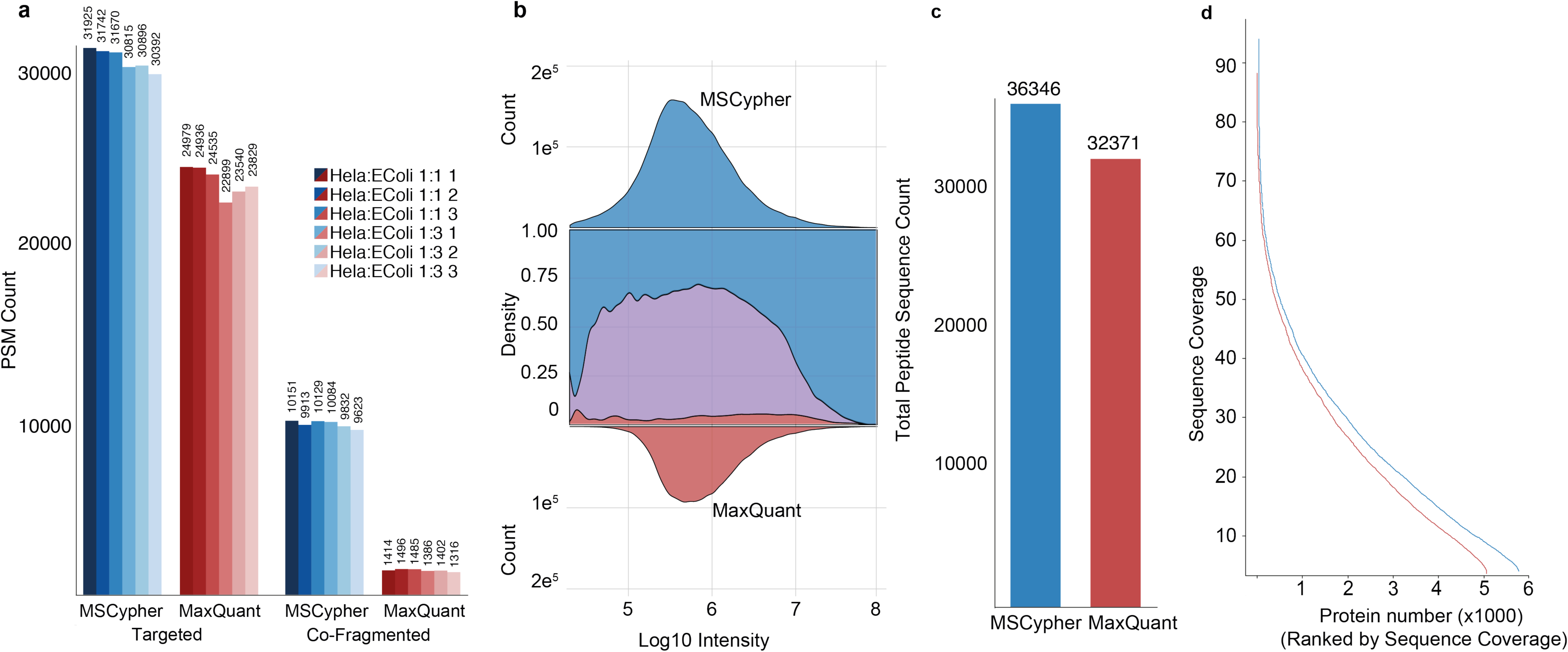
(a) Total count of PSM<u>s</u> per LC-MSMS run for both instrument picked (targeted) and co-fragmented precursors. (b) Top: frequency plot of MSCypher identified PSMs by log_10_ intensity. Middle: density plot of identified (blue) PSMs identified only by MSCypher, (red) PSMs identified only by MaxQuant and (pink/purple) PSMs identified by both workflows versus log_10_ intensity. Bottom: frequency plot of MaxQuant identified PSMs by log_10_ intensity. (c) Comparison of total peptide sequence count. (d) Protein sequence coverage by rank (highest to lowest coverage) where the red line represents proteins identified by MaxQuant and the blue line by MSCypher.

The count of unique peptides by file was improved by 30% over the MaxQuant search (**Figure. S3a**) and the level of missing values is lower than that observed by MaxQuant (**Figure S3b**).

To validate the accuracy of the 1% FDR we used two alternative means of checking for false positive identifications. Firstly, as the number of potential matches an isotopic cluster can obtain is not limited, the number of identifications assigned to a single feature (**Figure 4a)** represents an orthogonal measure of likely false positive assignments. Here, we show that less than 1% of the isotopic clusters detected have more than one identification assigned. Secondly, we performed a simple quantitative experiment that measured a quantitative FDR after seeding of *E. coli* tryptic peptides into HeLa tryptic peptides at two different concentrations (a ratio of 1:1 and 3:1 in triplicate **Figure 4b)**. **Figure 4c** shows a histogram of the log_2_ fold change that represents the three-fold increase in *E. coli* where <5% of the identified and quantified peptides fall beyond two standard deviations from the mean.

**Figure 4.**
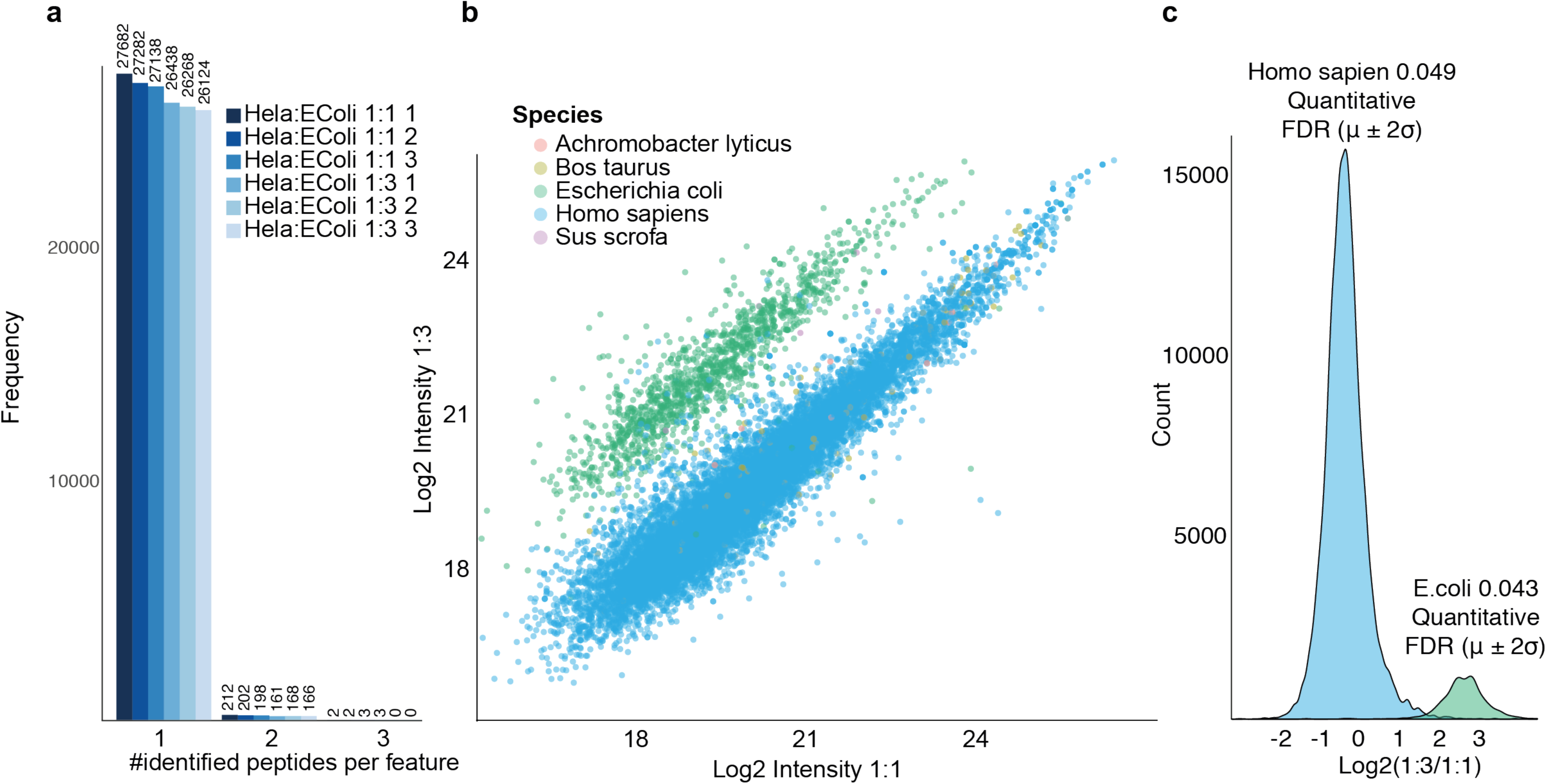
(a) Frequency of number of identifications assigned to a peptide feature. (b) Mean peptide intensity of triplicate runs for quantitative comparison of *E. coli* and human peptides, mixed at ratios 3:1 and 1:1. Only peptide features consistently identified in at least two replicates per group are shown. (c) Histogram showing the log_2_ ratio colored for species (blue = Homo sapiens, green = *E. coli*). Only peptide features consistently identified in at least two replicates per group are shown.

Future improvements in mass spectrometry technology that enhance dynamic range and sensitivity must consider the increased number of chimeric spectra that will be generated from complex samples. Thus, better strategies to maximize this natural multiplexing are needed to provide increased numbers of identifications of observable peptide features. Here we have introduced software that automatically fine tunes the spectra-peptide matching process using supervised machine learning and utilizes as much information as is available to describe a peptide spectral match, creating a larger separation power between true and false positive matches. Our approach is less sensitive to noise resulting from co-fragmentation of co-eluting peptides and yields both high sensitivity and specificity. We also include automated: retention time dependent intensity normalization to counter electrospray variation; label free quantitation (LFQ) pairwise comparison; and experimental evaluation and quality control plots (example reports for a published dataset (Choi et al., 2017) as **Supplementary HTMLs**). Overall, our workflow enhances the number of identifiable PSMs, peptide identifications, protein sequence coverage and is widely applicable to any form of proteomics and high-resolution mass spectrometer. In addition, this software provides detailed peptide- and fragment ion match statistics as a text output and will be crucial in order to further optimize fragment ion matching for different fragmentation techniques and acquisition strategies including data independent acquisition (DIA). MSCypher is freely available, an extensible workflow with detailed output that encourages further exploration of proteomics datasets via alternative machine and deep learning strategies (www.wehi.edu.au/people/andrew-webb/2372/resources-mscypher).

## CONTACT FOR REAGENT AND RESOURCE SHARING

Further information and requests for resources and reagents should be direct to and will be fulfilled by the Lead Contact, Andrew Webb.

## METHOD DETAILS

### Sample preparation for LC-MS/MS analysis

#### HeLa-ABRF spike-ins

Two sample sets containing lyophilised HeLa cell lysates (25 μg) spiked with four proteins at three different concentrations (20 fmol to 500 fmol) were provided by the ABRF 2016 PRG study. Sample 1 contained 100 fmol, Sample 2 contained no spiked in proteins, Sample 3 contained 20 fmol and Sample 4 contained 500 fmol. The spiked in peptides including ABRF-1 (beta-galactosidase from *E.coli)*, ABRF-2 (Lysozyme C, *G.gallus*), ABRF-3 (glucoamylase, *A.niger*) and ABRF-4 (Protein G, *Streptococcus spp.*). The protocol is largely based on the original sera-mag Speed bead (SP3) paper (Hughes et al., 2014), but with some differences. For all our experiments we used a 1:1 combination mix of the two types of commercially available carboxylate beads (Sera-Mag Speed beads, CAT# 90-981-121, 09-981-123, Thermo Fisher Scientific). Beads were prepared freshly each time by rinsing with water three times prior to use and stored at 4**°**C at a stock concentration of 20 μg/μl. DynaMag magnetic racks were used throughout the experiments (DynaMag-2, 12321D, Life Technologies). Spiked HeLa cell lysates were reduced with 500 mM Dithiothreitol (DTT, 50 mM final conc.) for 1 hr at 37 degrees **°**C. Samples were then alkylated with 500 mM Iodoacetamide (IAM) (100 mM final conc.) for 30 mins in the dark at room temperature (RT). Samples were quenched with 500 mM DTT (200 mM final conc.). Lysates were acidified with formic acid (pH <3) and a 2 μl of the concentrated bead stock (20 μg/μl) was added to samples. Acetonitrile (ACN) was then added to reach a final concentration of 50% (v/v). Mixtures were left to incubate upright at RT for 8 mins. Beads were then washed twice with 250 μl 70% ethanol and once with 250 μl acetonitrile. A further 100 μl acetonitrile was used to transfer the protein/bead suspension into a PCR plate for subsequent enzymatic digestion. Acetonitrile was completely evaporated from the PCR plate using a SpeedVac AES 1010 (Savant) prior to the addition of 10 μl digestion buffer (50 mM NH_4_HCO_3_) containing 1 μg Trypsin-gold (Promega, V5280). The plate was briefly sonicated in a water bath to disperse the beads, and the plate transferred to a PCR machine for overnight (∼16 hr) incubation at 37 **°**C. Following digestion, the PCR plate was lyophilised to dryness and the SP3 beads were then subjected to HILIC-based peptide fractionation. The beads were reconstituted in 100 μl 97% ACN, 5 mM Ammonium Formate (pH 12) and peptides were sequentially eluted off the beads using the magnetic rack. Six further fractions were collected (93% ACN, 89% ACN, 82% ACN, 75% ACN, 2% DMSO and 100 % ACN). Fractions were transferred to MS vials, lyophilised to dryness and reconstituted in 5 μl 0.1% formic acid/ 2% ACN.

#### HeLa-E.coli standard peptide mixture

A separate mixture of standard tryptic digests was created by mixing Pierce HeLa protein digest standard (#88328, Thermo Scientific) with MassPREP *E.coli* digestion standard (#186003196, Waters) at two different ratios (1:1 and 1:3).

### LC-MS/MS analysis

HILIC fractions (5 μl) and standard peptide mixtures (HeLa/E.coli) were separated by reverse-phase chromatography 1.9 μm C18 fused silica column (I.D. 75 μm, O.D. 360 μm x 25 cm length) packed into an emitter tip (IonOpticks, Australia), using a nano-flow HPLC (M-class, Waters). The HPLC was coupled to an Impact II UHR-QqTOF mass spectrometer (Bruker, Bremen, Germany) using a CaptiveSpray source and nanoBooster at 0.20 Bar using acetonitrile. Peptides were loaded directly onto the column at a constant flow rate of 400 nL/min with buffer A (99.9% Milli-Q water, 0.1% formic acid) and eluted with a 90 min linear gradient from 2 to 34% buffer B (99.9% acetonitrile, 0.1% formic acid). Mass spectra were acquired in a data-dependent manner including an automatic switch between MS and MS/MS scans using a 1.5 second duty cycle and 4 Hz MS1 spectra rate followed by MS/MS scans at 8-20 Hz dependent on precursor intensity for the remainder of the cycle. MS spectra were acquired between a mass range of 200–2000 m/z. Peptide fragmentation was performed using collision-induced dissociation (CID). The HeLa/E.coli standard peptide mixtures were run in triplicate.

### Data analysis workflow

The data analysis workflow consists of three steps: (1) Feature detection using MaxQuant, (2) Andromeda peak list file conversion to Mascot generic format, and (3) Integrated database searching and machine learning using MSCypher. The first step involves LC-MS feature detection, extraction, mass recalibration and identification of peptides through MaxQuant (Cox and Mann, 2008) and its built-in search engine, Andromeda (Cox et al., 2011). MaxQuant version 1.5.8.3 was used for all data analysis and the configuration files are provided as supplemental material. The second step involves converting the Andromeda peak lists (APL) and associated information for all features to Mascot generic format (MGF) using the APLtoMGFConverter (www.wehi.edu.au/people/andrew-webb/1298/apl-mgf-converter). The associated feature information (feature intensity, retention time apex as well as identified peptide sequence at 1% FDR are written to the MGF “TITLE=” line (see Figure S1) and used as the basis for “consensus” between search algorithms for retention time prediction and machine learning.

An example MGF “TITLE=” line appears as follows:

*TITLE=RawFile: Hela_Ecoli_1_1_20151124_1_01_3762 Index: 32937 Precursor: 0 _multi_ Charge: 2 FeatureIntensity: 2324000 Feature#: 2239887 RtApex: 2194.32 FeaturePif: 0.7024014 MS2Pif: 0.5880033 Ndp: 93 Ns: 25 Nip: 5 Seq: YLLGTSLAR Score: 51.346 #MS2: 1*

Converted MGF files are analysed using the MSCypher workflow. This includes several command-line programs which are instantiated *via* a configuration file (MSCypher_config.ini). MSCypher runs on Linux and Microsoft Windows 64bit operating systems utilising multiple cores/threads, as specified in the configuration file. External programs such as MSConvert, Elude and Percolator have wrappers in order to take advantage of the number of cores/threads (i.e. if there are 4 raw mass spectrometry files then all 4 will be simultaneously processed by these external programs if the number of “threads” is set to 4).

### MSCypher workflow

In addition to the steps detailed below there are a number of additional workflow options which can be specified in the configuration file depending on user requirements: (1) The MSConvert tool (ProteoWizard (Chambers et al., 2012; Kessner et al., 2008)) can be used for automatic generation of spectrum peak lists (MGF) from instrument specific files as an alternative to using MaxQuant; (2) The FastaSeqGenerator program can be used for generating a customised FASTA sequence database based on user-specified taxonomic identifiers. This program uses the MSPnr sequence database as input which is available from www.wehi.edu.au/people/andrew-webb/1295/andrew-webb-resources plus a list of taxonomic identifiers in a text file; and (3) An integrated process for *de novo* peptide sequencing using the PepNovo (Frank and Pevzner, 2005) program. If PepNovo is run then the sequencing results for all spectra are displayed alongside the database search results in the MSCypher output file.

Workflow options, input files, search parameters (e.g. modifications, enzyme cleavage rules), as well as output options are all specified in the MSCypher_config.ini file. The configuration file is automatically backed-up and time-stamped in the output folder for a specific analysis.

#### Database search

The MGF is searched with the Digger search algorithm. This step functions to identify all the potential peptide spectral matches (PSMs), and outputs 60 attributes for each spectrum. In addition to this, it generates decoys for every target sequence which are then used to score and rank potential matches. All candidate peptides (including user-selected potential modifications) that match a spectrum mass within a specified peptide tolerance (ppm, mmu or Da) are fragmented *in silico* and then matched and scored. Fragmentation is based on fragment ion types that are pre-defined and instrument-specific. The preliminary scoring stage was designed to accomplish both the elimination of low-scoring random matches and retention of correct assignments (if present), as well as the collection of ion-series matching spectral statistics for all target and decoy candidate peptides. The number of candidate peptides retained for each spectrum is currently 20 for both target and decoy. The preliminary score is based on an improvement of the “fast cross-correlation” calculation as implemented in Comet (Eng et al., 2015; Eng et al., 2013). In our method, the 15 most intense peaks per 110 Da bin are used in the matching process and the fragment ions that are generated are dependent on the amino acid composition of the peptide, fragment-ion type and precursor ion charge state. Currently 22 ion-series types are supported (immonium, a, b, b-98, b-18, b-17, b^2+^, b^2+^-98, b^2+^-18, b^2+^-17, c, c^2+^, y, y-98, y-18, y-17, y^2+^, y^2+^-98, y^2+^-18, y^2+^-17, YB, YA, z, z+1, z^2+^ and z+1^2+^).

Ion-series matching statistics for all target and decoy candidate peptides are collected during the preliminary scoring stage and used in the final score calculations. The number of candidate peptides matching a spectrum is recorded, with target and decoy matches recorded separately. For each candidate peptide, a set of theoretical fragment ions (which are dependent on both the amino acid composition and charge state of the precursor ion) are calculated and compared to the spectrum peak list. A “match” is recorded if the mass of the fragment ion matches (within the user-defined mass error) with peaks from a range within the 110 Da window. This range is defined as running from the least intense (Level 15) to the most intense (Level 1). For each “match” the following two pieces of information are retained: The matching peak’s intensity and intensity ranking (i.e. level 1-15), and the mass deviation of the “match” (i.e. mass difference between fragment ion and peak m/z) along with its fragment ion type (i.e. immonium or ‘y’ ion etc.). Multiple fragment ion matches to a peak are fractionally weighted according to their mass deviation. For example, if 3 different fragment ion types are all equidistant from, and match to, the same peak then they are awarded 1/3 of a “match”. Similarly, the “match intensity” is 1/3 of the peak intensity. The total number of matches to a particular ion type (stratified by level) is obtained by rounding-down the sum of the whole and fractional matches.

Additional information, in the form of matching statistics, is collected for all decoy candidate peptides. For scoring purposes, this constitutes the null model. When one considers the matrix of all candidate decoy peptides for each spectrum, each cell then contains a distribution of the number of peptides that match to fragment ions from a particular ion-series of a specified level. In order to combine information across ion-series (irrespective of type), the four highest “number of fragment ion matches” are recorded on a per-level basis.

As there are 20 peptides, at most, retained for each spectrum (based on preliminary scoring), the empirical decoy fragment ion statistics collected during this first scoring stage are used to re-score these peptides. The final scoring step involves the following processing steps for each spectrum: The number (N) of candidate decoy peptides matching the spectrum is determined; 2) The maximum number of fragment ion matches for each spectrum (M) is determined and stratified both by ion-series combinations (i, 1≤ i ≤ 4) and level (l, 1≤ l ≤ 15); and 3) The relative frequency of matching at least k out of M fragment ion matches is calculated and stratified by ion-series combination (i) and level (l). The ratio is log-transformed and converted to a tail score (S) (Eq. 1).

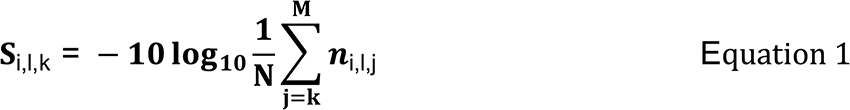

A linear least-squares regression of the tail scores for a particular cell (of a 4 x 15 matrix) allows for the calculation of both slope and intercept. A fitted line is extrapolated for target peptides that have a higher number of fragment ion matches when compared with all decoy candidate peptides for a given spectrum. All target and decoy candidate peptides that survive the preliminary scoring stage are then scored using the 4 x 15 matrix of slopes and intercepts. For each peptide, all ion-series are tested beginning with individual ion-series, and then subsequently testing combinations (up to a maximum of 4) for all ion-series. The score for a particular ion-series and level is therefore based on the number of fragment ion matches and is determined directly from the slope and intercept (“cell Score - cS” (see Eqn. 2)). The Peptide Score (Ps) is the maximum score after normalization, stratified by ion-series combinations and level. The normalisation (Int_MAf) is essentially a penalty for large mass errors on matching peaks and non-matching of high intensity peaks, and is used to break ties between peptides that have an identical number of fragment ion matches at a specific level. Int_MAf values are ≥ 1.0, because the total intensity of peaks at a particular level is divided by the mass error-weighted intensity of the matching peaks for that level.

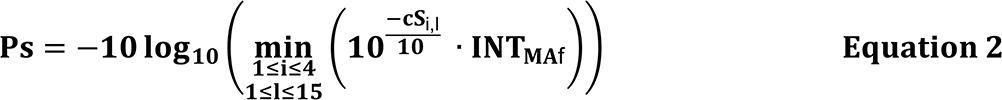

Subsequent changes to the original Digger search algorithm and scoring function (Kapp, 2013) are summarised below:

1) The fast cross-correlation (XCorr) score is calculated for all peptide-spectrum matches and serves as the preliminary score in retaining the top 20 target and decoy candidate peptides for each spectrum.

2) For each target candidate peptide, a decoy peptide is generated by retaining the n- and c-terminus residue and reversing the internal residues. A percent similarity (based on ‘b’ and ‘y’ ions) is then calculated between the target and decoy sequence taking into account all modifications. If the percent similarity between the target and decoy is greater than 20, then the internal residues are randomised rather than reversed. This concept, whilst similar to that implemented in Crux (McIlwain et al., 2014), represents an improvement over current implementations, since shorter peptides and peptides with multiple I/L or G amino acid residues negatively impact discriminating between target and decoy peptides.

3) Complementary ‘b’ and ‘y’ ion calculations are performed for all peptides taking into account all modifications in order to calculate the ‘percent bond cleavage’ i.e. percentage of backbone bonds cleaved for a particular peptide based on the complementary n- and c-terminal fragment ions. This is an important feature for the machine learning classification step since the ideal “correct” hit would be represented by a ladder of sequence ions confirming n-1 amino acid residues.

4) For each MS2 peak list the number of selected “most intense” peaks used for scoring purposes was previously 10 per 100 Da window. This has been increased to 15 peaks per 110 Da windows to enable the detection of lower abundant fragment ions derived from co-isolated precursor ions in MS1 (i.e. chimeric spectra).

5) Options are provided in the configuration file that allows finer control over whether the intensity of peaks are used for penalising the Digger score for a particular PSM. For example, if two peptides match the same number of peaks, but the intensities of the matched peaks differ, then the Digger score would reflect this situation if the “IntensityPenalty” was “on”. For the current analysis, the “IntensityPenalty” was set to “off” since the machine learning step uses all 60 features in training and classification.

Digger outputs a tab-delimited text file containing columns describing all the spectrum matching attributes (see Table S1) for all potential PSMs (includes rank 1-20 target/decoy matches), which is the master table that is used as input to all the subsequent processes.

#### Retention time prediction

Elude (Moruz et al., 2010) was used to predict the retention times of all potential PSMs utilizing a training set, which selects from a pool of consensus matches (filtered from Andromeda and Digger concurring on a specific PSM) for each mass spectrometry run. The selection process is designed to include peptides across the retention time of all runs in order to capture all amino acid properties, including modified peptides (e.g. phosphorylation) that ultimately affect the elution times. All consensus modified peptides are included in the training model, whilst a selection process is enabled for all unmodified peptides. The highest intensity consensus unmodified peptide is selected per 3 secs windows across the whole MS run. If Andromeda is not used, then consensus is based on a combination of the Digger probability score (>45), XCorr (>2.5), deltaCn (>0.1) and peptide rank (i.e. peptide is rank 1 by Digger probability score and Xcorr).

#### Machine learning

The 60 attributes (features) that characterize a PSM (Supplementary Table S1) are extracted from the MSCypher text file. Importantly, attributes such as Digger score or delta scores (score difference rank 1-5) are not used in the machine learning step since these attributes could bias model training because of the dependent target-decoy competition during the database search. The top 5 ranked target and decoy matches for any given MS/MS spectrum are extracted and labelled as “known-set” of PSMs to be any match to the target database where there is consensus between search algorithms (total of 139331 PSMs) and any match to the decoy database (total of 8193480 PSMs). The main parameters of the Random Forest (RF) were set as follows: the number of trees was 5000, to reduce variation between models and increase separation between decoy and target matches, the sample size was set to the square root of the number of target database matches used in the training set, to assure the model is built using balanced classes; all the other parameters were left at default, with the exception of “nodesize”, for which 4 different values (5,10, 20 and 30) were tested, to investigate its effect on computational requirements for generating the model.

To assess the accuracy of the model for varying parameters and training set size, we picked a varying number of target PSMs (2, 5, 10, 30 thousand) at random from the “known-set” pool, and a varying number of decoy database matches as a factor of target PSMs (1x, 5x, 10x). Figure S2 shows the results for model accuracy, false positive rate (FPR) and true positive rate (TPR), using the remainder of the “known-set” for validation, with different training set sizes and values for the node-size parameter, and by repeating each estimation ten times. Also, the figure shows the time required for generating the model. The accuracy of all the models was above 99.5%, however a 0.2% variation from the lowest to the highest accuracy can account for a very large number of PSMs when analyzing large datasets. Thus, we used a training set of 180K PSMs with a target to decoy PSMs ratio of 1:5 as a standard training set and the “*nodesize”* parameter set to 5 for all the datasets for this work. This combination showed a balance between the highest accuracy and TPR with the lowest FPR and the time spent for generating the model by parallelizing the computation on four cores.

After generating the model, it was used to predict targets on the complete dataset. The output of the RF is represented as a vote count, where each tree of the forest accounts for one vote by classifying a PSM as target or decoy match, we take the sum of the votes that classify a potential PSM as a decoy database match as the preliminary score of a PSM. Supplemental Figure 3 shows the density distribution of scores for PSMs from both the target and decoy matches derived from the random forest algorithm. In this case, the scores range from 5000 to 0, where each vote accounts for the output of a tree in the forest where the tree classifies the PSM as a decoy DB match. The figure shows the wide separation between true target database matches (on the right side with low to 0 RF votes) and random decoy matches (on the left side with high to 5000 RF votes) that this classification approach is capable to generate, thus highly increasing the ability to recognise a potentially true spectrum match.

From the number of RF votes that classify the PSM as a decoy match, we calculated a PSM- score as follows

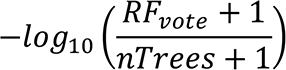

Taking the top-ranking spectrum match for each PSM, we group them by sequence length (6 to 25) and group together all sequences with a length greater than 25. We then applied a 1% FDR filter to each group independently, experiment wide.

As a second FDR cut-off filter we calculate the score for a modified peptide feature (intended as the combination of an identified peptide’s sequence, modification and charge state) by calculating its maximum PSM score experiment wide and then filtering at 1% FDR.

Machine learning using Percolator to calculate q-values and PEP scores for all peptides is included in the workflow and reported in the MSCypher output table if this option is enabled in the MSCypher configuration file. Turning this option on by default is useful as it provides additional evidence and confirmation of the reported significant PSMs.

#### Protein inference

Protein inference was performed using only unique significant scoring peptide features from the Random Forest output, by summing the peptide feature scores for all target and decoy inferred proteins. Proteins are classified into six hierarchical categories: distinct, differentiable, subsumable, superset, subset and equivalent in accordance with DBParser (Yang et al., 2004) guidelines and subsequent refinement (Huang et al., 2012). MSCypher only reports proteins from the distinct, differentiable and superset categories adhering strictly to the Occam’s razor principle to ensure that the minimum set of proteins are reported at 1% FDR that explain all unique significant scoring peptide features. Target and decoy protein lists including identified peptides and scores are output including a protein FDR table indicating the number of protein groups at different FDR percentages.

#### Label-free quantitation and statistical analysis

A label-free quantitation workflow written in R automatically reads the protein inference tables, experiment design (experiment_design.txt) and a table that lists the pairwise comparisons to be performed (pairwise_comparisons.txt). This is a modified version of the workflow previously reported by our group (Delconte et al., 2016). In the script, a feature was defined as the combination of peptide sequence, charge and modification. For each pairwise comparison, features not found in at least half the number of replicates in at least one of the conditions were ignored.

Intensity normalisation was performed by selecting a LC-MS/MS run with the highest number of identified features as a reference and removing non-linear variation of intensities with respect to retention time by performing a Loess regression (Cleveland, 1992) on matched features. Features assigned to the same protein differ in the range of intensity due to their physio-chemical properties and charge state, to further correct for these differences, each intensity value was multiplied by the ratio of the maximum of the median intensities of all features for a protein over the median intensity of the feature.

Missing values where imputed using a random normal distribution of values with the mean set at mean of the measured distribution of intensity values minus 1.8 standard deviations (s.d), and a s.d. of 0.3 times the s.d. of the distribution of the measured intensities (Cox et al., 2014). The probability of differential expression between groups was calculated using the Wilcoxon Rank Sum test excluding any non-unique sequences, the output of the R function wilcox.test included P value, confidence interval and ratio estimate. Probability values were corrected for multiple testing using Benjamini–Hochberg method. Cut-off lines with the function y = −log_10_(0.05) + c/(x - x_0_) were introduced to identify significantly enriched proteins (Keilhauer et al., 2015). “c” was set to 0.2 while “x_0_“ was set to the standard deviation of the protein log_2_ ratio. The script outputs a results table (LFQ_Results.txt) with all the information on the pairwise comparisons at the feature level. A protein table (LFQ_Proteins.txt) with protein log_2_ ratios, PValues, confidence intervals using both Wilcoxon rank sum test and Student’s t-test by pairwise comparisons, peptide counts and sum peptide intensities for the whole experiment by group and by LC-MS/MS run. A peptide table (LFQ_Peptides.txt) similar to the protein table, but with quantitative analysis at the peptide level. Lastly an HTML document is provided that contains quality control plots to evaluate the experiment (see **“Supplemental_LFQ QC Report.html”** file). Additionally, an interactive volcano plots HTML file is generated for protein quantitative analysis for each pairwise comparison (see **“Supplemental_LFQ Results.html”** file).

#### Visualisation and support for external software tools

Output files that are compatible with 3^rd^ party software tools are supported as part of the workflow. A text file containing protein accession numbers and peptide sequences are written out (Protter_input.txt) which can be loaded into the Protter on-line tool (Omasits et al., 2014)(http://wlab.ethz.ch/protter/start/). A “peptigram” csv file is output summarising peptide feature intensities across all mass spectrometry runs, conditions and bioreplicates for all inferred proteins. This file can be used as input to the on-line Peptigram program (Manguy et al., 2017) (http://bioware.ucd.ie/peptigram/) for visualisation. A Skyline (MacLean et al., 2010) file (Skyline_input.ssl) and protein sequence FASTA file for all inferred protein hits for the analysis are written out, which can be loaded directly into Skyline in order to build libraries, visualise annotated spectra and enable verification and follow up for targeted experiments. Fully compatible MSstats (Choi et al., 2014) peptide and protein input files are written out which can be loaded and run automatically in R with minor manipulation. Finally, the program Cd-hit (Li and Godzik, 2006) is run as part of the workflow in order to cluster protein sequences at 90% and 60% based on the list of inferred proteins. These files could be used as input to a follow-up/cascaded search or error-tolerant type database searches.

### Comparison of multiple search algorithms

The Human/E.coli dataset was analysed using 4 different search algorithms (MaxQuant/Andromeda, MSCypher/Digger, MSGF+ and Comet) using identical input, search parameters and the *Homo sapiens* / *Escherichia coli* plus contaminants protein sequence database. Variable modifications specified were N-term protein acetylation, methionine oxidation and N-term pyro-glutamic acid and Cys+57 as a static modification. Precursor and product ion mass tolerances were 10 and 20 ppm respectively, except for Mascot where the product ion mass tolerance was set at 20 mmu. Default search parameters appropriate for the data were used for Comet and MSGF+ with the aim to minimise differences in the search space and maximise consensus between the different search algorithms. The search results for all 4 algorithms were then combined in order to compile a large set of consensus spectra for generating “chimeric” spectra using Monte Carlo simulations.

### Monte Carlo simulation of chimeric spectra

There were 770000 consensus PSMs from the Human/Ecoli dataset. Using a Monte Carlo simulation, 110 Million PSMs were generated in total by randomly selecting 100 spectra from the consensus set and simulating this process 10000 times (simulations). This first set represented iteration 0 (i.e., original spectra). For iteration 1 this process was repeated, but an additional randomly selected peak list was seeded in i.e., all peak lists generated were of increasing complexity mimicking semi-real chimeric spectra. This process was repeated 10 times so that iteration 10 peak lists would be the most complex and mimic wide isolation windows not normally utilised in DDA shotgun proteomics experiments, but routine in DIA (e.g. variable SWATH windows). No attempt was made to seed in peak lists of differing complexity or intensity. The 110 Million PSMs were batched into MGF files and named accordingly with the simulation and iteration recorded in the file name (e.g. monteMGF_s4000_i7.mgf). These 110000 input files were then searched against MSCypher/Digger, MSGF+ and Comet search algorithms. The results for the 3 search algorithms were then summarised based on the original consensus peptide sequence which is recorded in the MGF Title line for each spectrum.

### Software availability

The MSCypher workflow binaries for Windows and Linux, detailed instructions, example datasets are freely available from www.wehi.edu.au/people/andrew-webb/2372/resources-mscypher. Source code will be available from Github

(https://github.com/MSCypher).

## Cell Systems

### Supplemental Information

|  |  |
| --- | --- |
| Figure S1 | E.coli human APLToMGF Conversion parameters |
| Figure S2 | Machine learning model evaluation |
| Figure S3 | Random Forest vote count distribution |
| Table S1 | MSCypher tab-delimited output file contents |

**Supplementary Figure S1.**
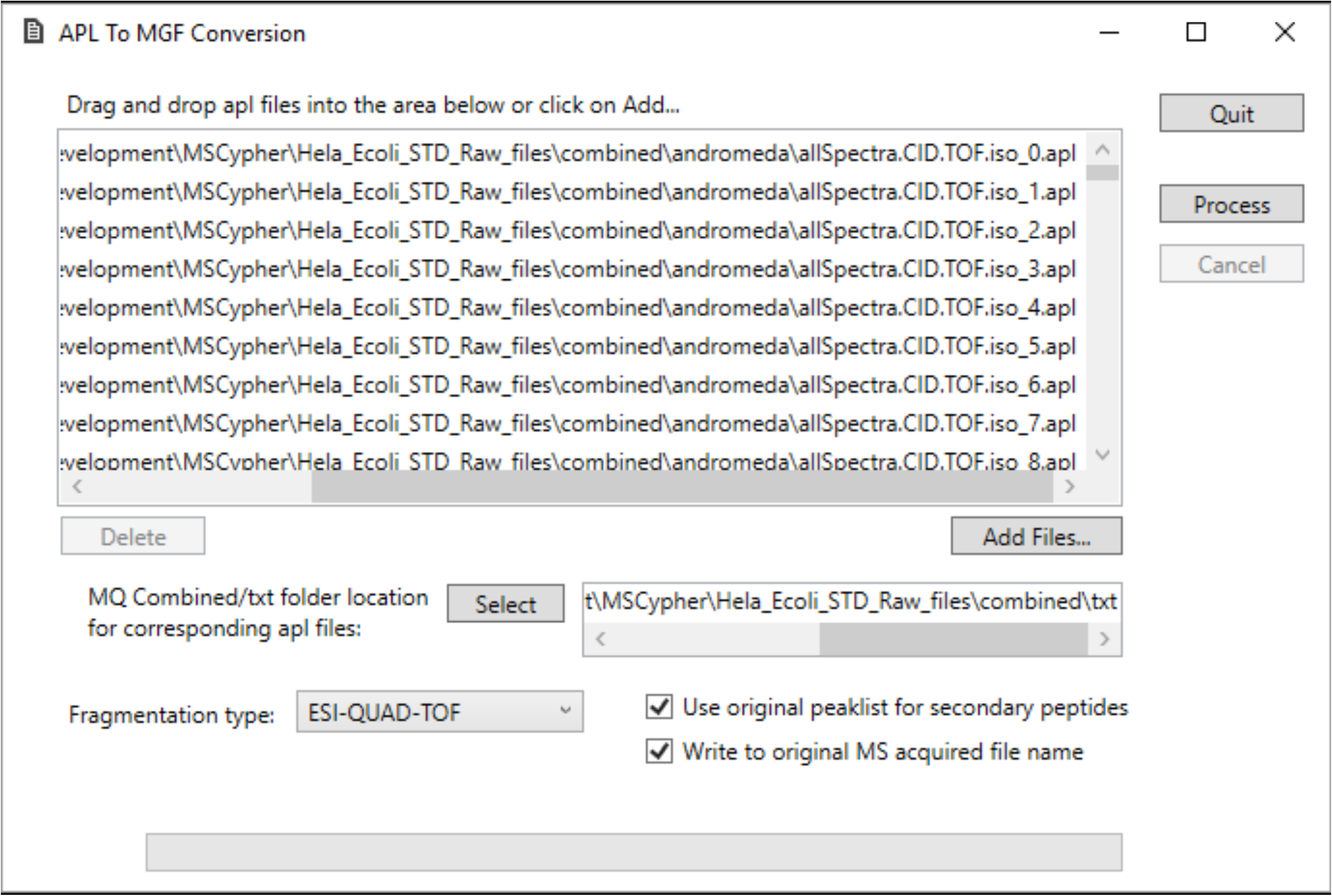
APLToMGFConverter graphical interface showing file and parameter settings for converting MaxQuant/Andromeda peak lists (APL) to Mascot generic format (MGF) for MSCypher analysis. The original MS2 peaklist was used for co-isolated and co-fragmented peptides (i.e. secondary peptides).

**Supplementary Figure S2.**
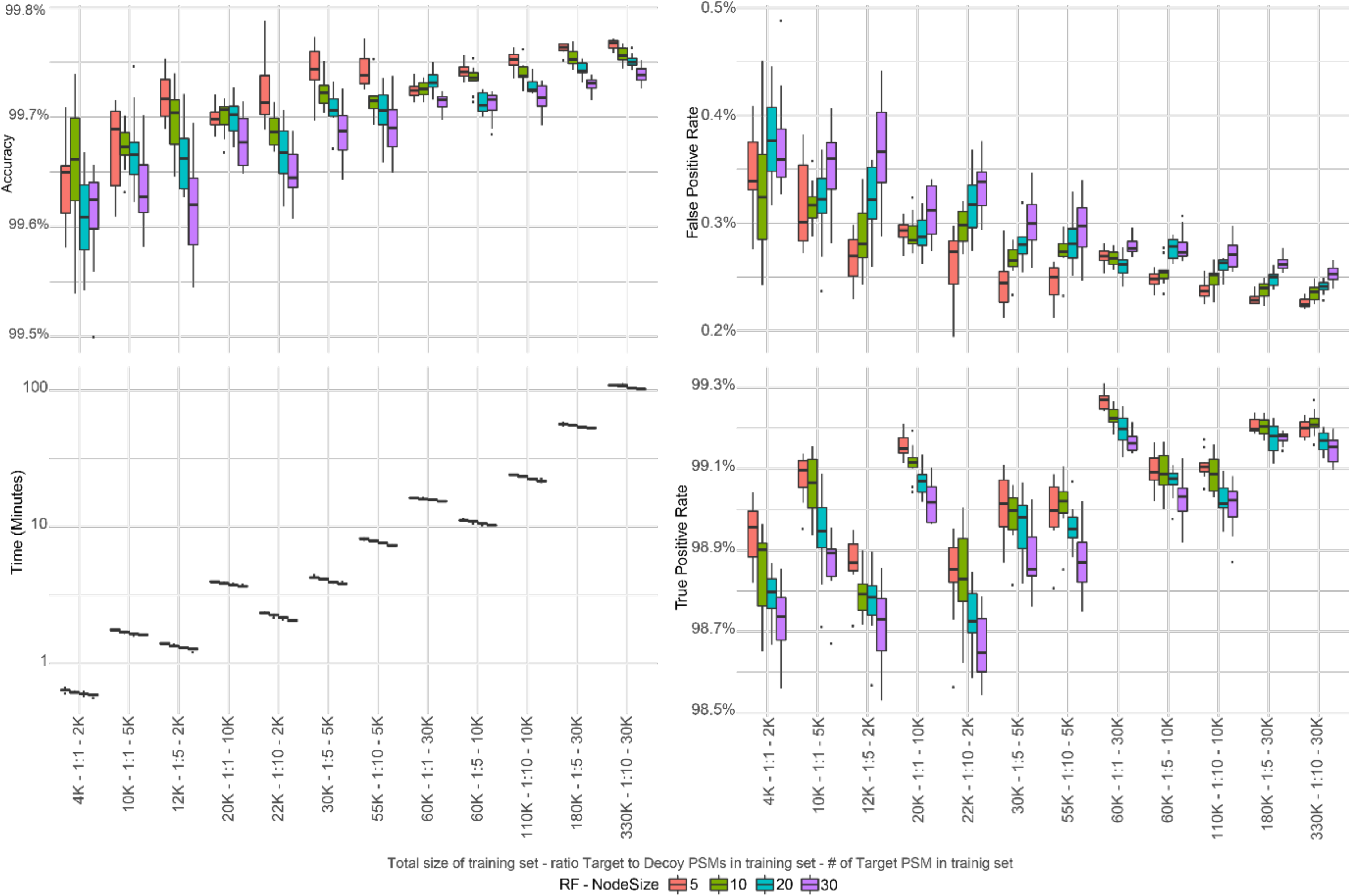
Accuracy (top-left), false positive rate (top-right), true positive rate (bottom-right) and computing time (bottom-left) of multiple random forest models using varying training set size and *nodesize* parameters. Each combination was assessed ten times by taking a different random set of PSMs to assess model variance at varying training sets.

**Supplementary Figure S3.**
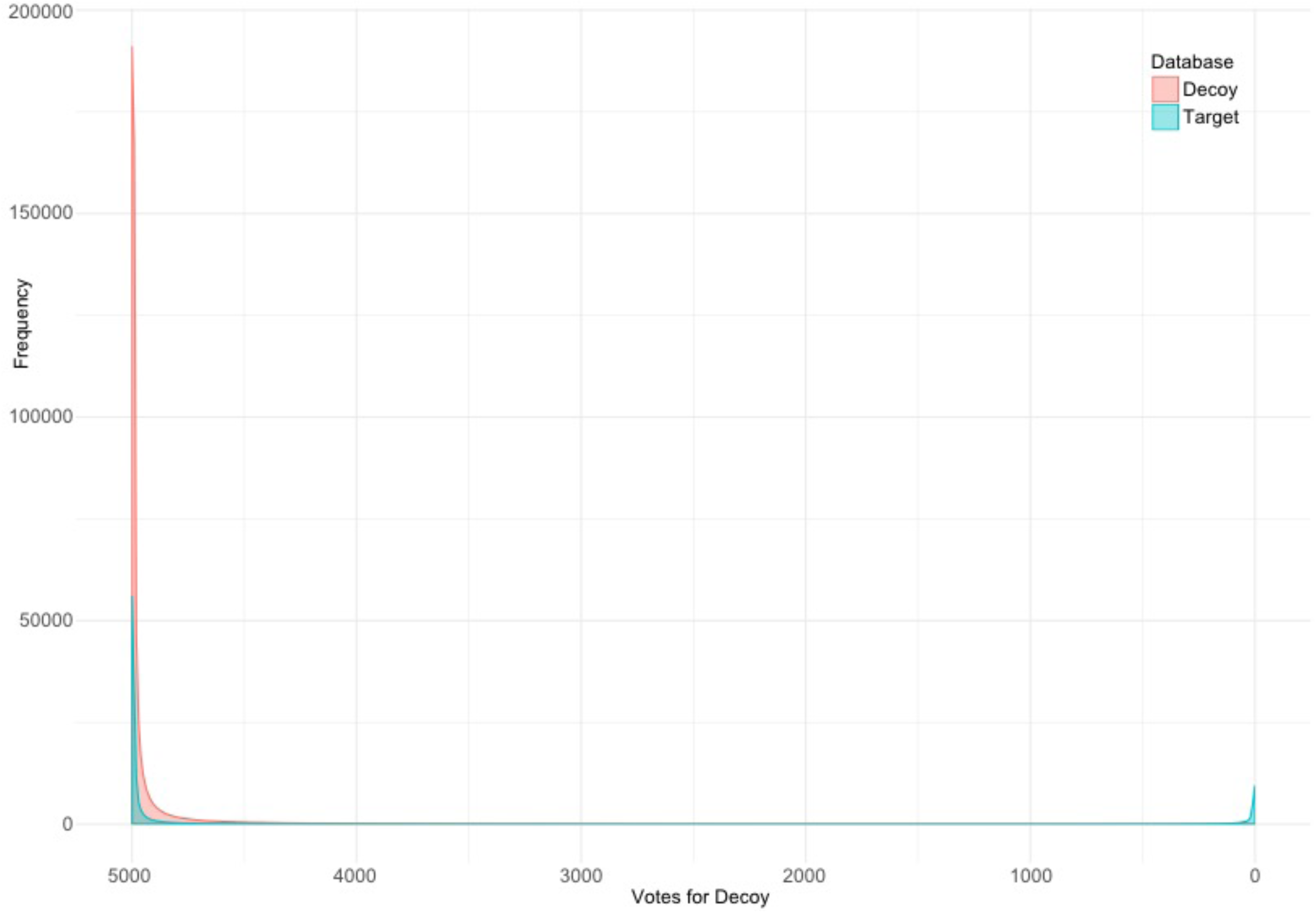
Distribution density plot of the rank one potential matches from Target (red) and Decoy (blue) databases.

**Supplementary Table S1.** MSCypher tab-delimited output file detailing all PSM match information used throughout the workflow by all processes.

| Column name | Description |
| --- | --- |
| QueryNum | Query number used internally per input file (unique to an input file) |
| Target | Target or Decoy indicating whether the peptide sequence is a real entry from the sequence database, reversed or semi-randomised |
| Rank | Ranking of the peptide spectral match based on the Digger score (1-20) |
| NumPrecMatches | Number of candidate peptides (target only) that match the spectrum based on searched mass and tolerance |
| PepIDConsensus | Value between 0-1000 that reflects consensus and scoring properties with 0 indicating a poor match and 1000 a perfect match |
| PepPIF | Precursor ion fraction (MS1) derived from MaxQuant output |
| MS2PIF | Precursor ion fraction (MS2) derived from MaxQuant output |
| NumScoringFragIonMatches | Number of fragment ion matches used in deriving the Digger score |
| NumScoringFragIonPeaks | Number of spectral peaks used for the matching fragment ions in deriving the Digger score |
| PercentSimilarity | % Similarity between target (real) and decoy peptide sequence based on 'b' and 'y' ions taking into account modifications |
| PepSeq | Actual peptide sequence identified from the sequence database |
| deNovoSeq | deNovo sequence derived from the spectrum using the PepNovo program |
| rnkScr | PepNovo rank score |
| pnvScr | PepNovo Pevsner score |
| EludePepSeq | Modified peptide sequence using Unimod representation (e.g. M[unimod:35]MEFK) |
| VMLScore | Variable Mod Localisation probability score (e.g. GHFS(1.0)DGS(0.1)LK) |
| Mods | Actual modification mass and site (e.g. 15.994915@1) |
| NumMods | Number of observed variable modifications |
| NumPhosphos | Number of phosphorylation modifications |
| PhosphoMods | Same as "Mods" but specific to phosphorylation only modifications |
| DiggerPepScore | Digger search algorithm probability based score |
| DiggerPepHomologyScore | Concept of a homology "gap" score (i.e. gap score plus 13) |
| PercolatorQValue | Percolator's q-value (i.e. global FDR) |
| PercolatorPEP | Percolator's PEP score (i.e. posterior error probability) |
| FeatureIntensityAUC | Peptide or feature MS1 intensity based on area under curve of all isotopes |
| ObsRtMS2 | Observed MS2 retention time of peptide or feature in minutes |
| ObsRtApex | Observed retention time Apex of peptide or feature in minutes |
| PredRt | Predicted retention time (using Elude) of peptide or feature in minutes |
| MZ | Observed precursor m/z value (from MS1 scan after recalibration if any) |
| Charge | Charge state of precursor ion |
| ExpMassDa | Observed neutral mass of feature (Mr) in Daltons |
| PepMassDa | Calculated neutral mass of peptide (Mr) in Daltons |
| DeltaMassDa | Difference in calculated and observed mass of peptide in Daltons |
| DeltaMassPPM | Difference in calculated and observed mass of peptide in parts-per-million |
| C13 | Values between 0-2 – e.g. if C13 is 0 then the monoisotopic peak has been matched |
| NumDataPoints | Only valid if the data has been processed by MaxQuant feature detection (from allPeptides file) |
| NumScans | Only valid if the data has been processed by MaxQuant feature detection (from allPeptides file) |
| NumIsotopePeaks | Only valid if the data has been processed by MaxQuant feature detection (from allPeptides file) |
| FileName | Name of the input file |
| MS2Scan | Instrument MS2 scan number for peptide spectral match |
| RMSErrorPPM | Root mean square error taking into account all fragment ion matches in parts per million |
| RMSErrorNumMatches | Number of fragment ions used in the RMSErrorPPM calculation |
| TIC | Total Ion Current for MS2 spectrum |
| CompBondCleavage | Percentage complementary bond cleavage i.e. 100% would mean that every backbone bond is broken into complementary b and y matching fragment ions |
| DeltaScore | Difference in score between rank X hit and rank 1 hit |
| DeltaRt | Difference between observed and predicted retention time in minutes |
| ChimericStatus | True if the feature is targeted by the instrument (i.e. precursor) or False if the peptide was co-isolated in the isolation window (i.e. secondary) |
| FeatureNum | Recorded feature number used internally – also represents the line number in the AllPeptides text file from MaxQuant |
| MissedCleavages | Number of missed cleavages if an enzymatic search |
| RPMS | Relative proton mobility scale – for future use |
| SVMScore | Support vector machine score – alternative to Percolator machine learning called Newbee |
| SVMSig | Support vector machine score is significant – true or false (Newbee) |
| SVMResult | Support vector machine prediction – target or decoy (Newbee) |
| RFFeatureScore | RandomForest feature score |
| RFPSMScore | RandomForest peptide spectral match score |
| ModPepID | Internal index for tracking peptides based on their modification status |
| Rawfilename | Original acquired raw mass spec filename |
| RFRank | RandomForest ranking of peptide spectral match |
| RFFeatureFDR | RandomForest false discovery rate at the feature level |
| RFPsmFDR | RandomForest false discovery rate at the peptide spectral match level |
| TargetVotes | RandomForest number of votes awarded (max 5000) for target hit |
| DecoyVotes | RandomForest number of votes awarded (max 5000) for decoy hit |
| RFPrediction | RandomForest prediction i.e. either target or decoy |
| MAErrorPPM | Fragment ion mass accuracy mean error in parts-per-million |
| AltDeltaScore | Digger delta score |
| FragMZ | List of matching fragment ion m/z values (semi-colon separated) |
| FragInt | List of matching fragment ion intensity values (semi-colon separated) |
| FragError | List of matching fragment ion delta mass values in daltons (semi-colon separated) |
| FragIonTypes | List of matching fragment ion types (semi-colon separated) |
| FragPos | List of matching fragment ion positions in peptide sequence (semi-colon separated) |
| FragCharge | List of matching fragment ion charge values (semi-colon separated) |
| Consecutive ion reward | Cumulative reward for matching consecutive fragment ions (22 columns) |
| Summed intensity | Summed intensity of matching fragment ions for 22 ion-types (22 columns) |
| Number of matching ions | Count of matching fragment ions for 22 ion-types (22 columns) |
| StartPos | Start position of peptide in protein sequence (semi-colon separated for multiple proteins) |
| EndPos | End position of peptide in protein sequence (semi-colon separated for multiple proteins) |
| PreRes | 3 residues preceding the peptide for its associated protein (semi-colon separated for multiple proteins) |
| PostRes | 3 residues after the peptide for its associated protein (semi-colon separated for multiple proteins) |
| EnzymeSpecificity | Complete, semi or non-specific (2,1 or 0) |
| ProteinLength | Length of protein (semi-colon separated for multiple proteins) |
| AccNums | List of protein accession numbers (semi-colon separated) |
| FastaEntryNum | Actual entry number in FASTA file (zero-based index) |
| UniqueID | Index into spectral peaklist file (e.g. MGF) for other applications e.g. Skyline |
| deltaCn | Delta cross-correlation score reflecting peptide ranking for MS2 peaklist |
| XCorr | Cross-correlation score between peptide and MS2 peaklist (i.e. sequest xcorr) |

